# In vivo Biomechanical Assessment of Iridial Deformations and Muscle Contractions in Human Eyes

**DOI:** 10.1101/2022.01.14.476393

**Authors:** Babak N. Safa, Mohammad Reza Bahrani Fard, C. Ross Ethier

## Abstract

The iris is a muscular organ whose deformations can cause primary angle-closure glaucoma (PACG), a leading cause of blindness. PACG risk assessment does not consider iridial biomechanical factors, despite their expected influence on iris deformations. Here we exploited an existing biometric data set consisting of near-infrared movies acquired during the pupillary light reflex (PLR) as a unique resource to study iris biomechanics. The PLR caused significant (>100%) and essentially spatially uniform radial strains in the iris *in vivo*, consistent with previous findings. Inverse finite element modeling showed that sphincter muscle tractions were c. 5-fold greater than iridial stroma stiffness (range 4- to 13-fold, depending on sphincter muscle size). This muscle traction is greater than has been previously estimated, which may be due to methodological differences and/or to different patient populations in our study (European descent) vs. previous studies (Asian); the latter possibility is of particular interest due to differential incidence rates of PACG in these populations. Our methodology is fast and inexpensive and may be a useful tool in understanding biomechanical factors contributing to PACG.

## Introduction

The human iris is an annular tissue disc with remarkable properties, including extreme contractility, e.g., iridial contraction can cause pupil diameter to change from 1 to 9 mm in a fraction of a second (Newsome and Loewenfeld 1971). Furthermore, the iris’s contractions and its anatomical placement in the anterior chamber (Figure 1A) involve the iris in glaucoma, the leading cause of irreversible blindness worldwide (World Health Organization 2019). Specifically, in the common form of glaucoma known as primary angle-closure glaucoma (PACG), the iris impedes aqueous humor drainage from the eye, drastically elevating intraocular pressure (IOP) and leading to a potentially blinding medical emergency (Friedman et al. 2012).

**Figure 1:**
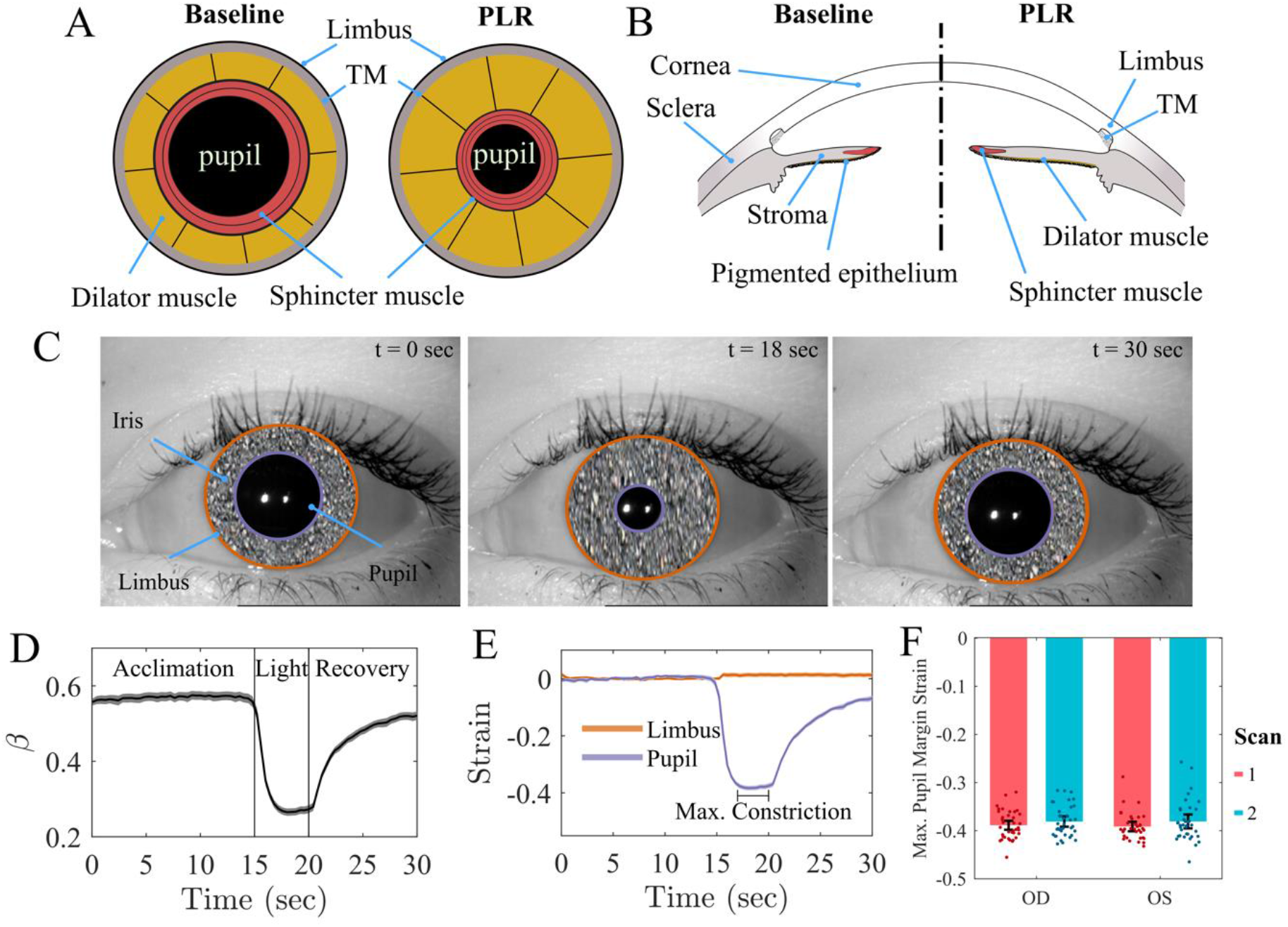
Schematic illustration of the mechanics of pupillary light reflex (PLR) in frontal **(A)** and **(B)** sagittal views, showing the pupil, the iris, and its attachment to the limbus and trabecular meshwork (TM). When the circumferential sphincter smooth muscle is activated, the pupil constricts (i.e., PLR). **(C)** Three representative images of the PLR from the same subject at the beginning of the test (t = 0 sec), during maximum constriction (t = 18 sec), and at the end of recovery (t = 30 sec). We have obscured iridial surface features to protect the identity of the subject. **(D)** The ratio of the pupil radius to limbal radius (***β*** = ***r***_***p***_/***r***_***l***_; mean as solid line and 95% confidence interval [CI] as shaded area). Initially, the pupil accounted for 56.6% ± 7.0% (mean ± std deviation) of the iris diameter (limbus edge diameter), while at maximum pupil constriction, it reduced to 26.2% ± 4.3%. For purposes of these calculations, we averaged the test-retest measurements for each eye. **(E)** Throughout the PLR test, the limbus diameter did not change and had a negligible strain (1.3% ± 4.3%; single-group t-test compared to zero p=0.007). After light exposure, the pupil demonstrated a dramatic 38.6% ± 3.1% (single-group t-test compared to zero p<0.001) compressive strain. The graph shows mean & 95% CI over all subjects (shaded area, difficult to distinguish because it is small). **(F)** The tested subjects’ peak pupillary margin strain at PLR was not different between the left (OS) or right (OD) eyes (p=0.827), and the results were repeatable between scans (p=0.099). Here, individual data points are shown overlaid with the error bars indicating 95% CI.

Risk factors for PACG include anatomical deficits (e.g., a crowded anterior chamber), age, and genetic background (Friedman et al. 2012); however, these factors alone cannot predict PACG incidence. For instance, the large ZAP trial (He et al. 2019) showed that only a small percentage of people classified as high-risk progress to PACG within six years, indicating that currently accepted risk factors are incomplete and inadequate.

Iris biomechanics, which strongly influences iridial deformations, is likely to be an additional risk factor for PACG. For example, dilation of the pupil induces a concave curvature of the iris favorable for developing PACG (Amini et al. 2012). Importantly, patients with a history of PACG tend to have an iris with higher stiffness and lower permeability (Pant et al. 2018; Panda et al. 2021), suggesting clinical utility in knowledge of *in vivo* iridial biomechanical properties. However, specific biomechanical risk metrics for PACG remain unknown, in part due to the difficulty of characterizing *in vivo* mechanical proprieties of the iris

Although the iris is optically accessible, its structure is complex, posing challenges to understanding its biomechanics and structure-function relationships. Notably, iridial contractions are driven by two antagonistic smooth muscles, i.e., the sphincter and dilator muscles (Figures 1A and B). Their contractions change iridial morphology (e.g., iris volume (Quigley et al. 2008)), mechanical properties (e.g., stiffness (Whitcomb et al. 2009) and permeability (Tan et al. 2019)).

Here we evaluated the *in vivo* biomechanics of the iris, exploiting the fact that iridial deformation is of interest in a wide range of scientific and technological applications (Pamplona, Oliveira, and Baranoski 2009). Specifically, because iris surface features are unique to each individual and are stable throughout life (Daugman 2004), iris recognition is widely used in biometric identification, which has motivated the acquisition of large data sets containing movies of human iridial motion during the pupillary light reflex (PLR) (Omelina et al. 2021). We used one such publicly available biometric data set, consisting of near-infrared (NiR) videos of human irides during PLR, which allowed us to calculate *in vivo* iridial strains and estimate muscle traction. We observed strains of larger than 100% and muscle tractions 5-fold greater than iris stromal stiffness. The methodology described herein provides a novel approach for *in vivo* evaluation of iris biomechanics using an accessible imaging modality, thus laying the groundwork for future clinical and functional assessment of iris biomechanics in the pathophysiology of glaucoma.

## Results

### Pupil and limbus deformations during PLR

The iris is highly sensitive to light, with the pupil constricting in response to an increase in light intensity during the PLR. To biomechanically analyze the iris, we quantified iridial deformations by tracking the limbus and pupil during PLR (Figure 1A-C), using images from the publicly available *Warsaw-BioBase-Pupil-Dynamics v3* data set (Kinnison et al. 2019). We used digital image segmentation and Daugman’s method (Daugman 2004; Sivaraman 2021) to calculate the limbal and pupillary diameters throughout 30-second videos (n=163 videos from 42 subjects; Figure 1C) and calculated the ratio of pupillary to limbal radii (*β* = *r*_*p*_/*r*_*l*_) and hence the Lagrangian strains (*ϵ*_*θθ*_) of the limbus and pupil margins. As expected, the limbus did not appreciably deform during PLR (Figure 1C and D), and the pupil maintained a constant radius in darkness (dark adaptation during acclimation phase; Figure 1D). However, the pupil dramatically contracted when the eye was exposed to ambient light (Figure 1D and E), gradually returning towards baseline after light stimulation ended (Figure 1D and E). The average *β* during acclimation was 56.6% ± 7.0% (mean ± std. deviation), reducing to 26.2% ± 4.3% at maximum constriction (Figure 1D). The average strain of the limbus margin was negligible (*ϵ*_*θθ*_ = 1.3% ± 4.3%; single-group t-test compared to zero, p = 0.007), while at maximum pupil constriction the pupillary margin strain was *ϵ*_*θθ*_ = −38.6% ± 3.1% (p < 0.001; Figure 1E).

We next analyzed variability in pupillary margin strain between scans for the same eye (test-retest) and between fellow eyes from the same subject. A difference in the PLR between the left (OS) and right (OD) eyes is known as a relative afferent pupillary defect, RAPD, and can indicate an underlying medical condition (Chang et al. 2013). However, we saw no evidence of RAPD in the 42 pairs of eyes in the data set (p = 0.977, linear mixed-effects model [LME]), and the test-retest paradigm did not result in different PLR responses (p = 0.029; Figure 1F). Although each eye’s test-retest scans indicated that the pupillary margin strain was slightly smaller in the second scan, the size of this effect was small, with less than 1% strain difference (0.8% ± 2.9%), which indicates that the PLR provides repeatable metrics.

### Spatial distribution of mechanical strain in the iris

Although deformation at the pupillary margin is of interest, more information can be obtained by determining local deformation across the iris stroma. We therefore performed digital image correlation (DIC (Palanca, Tozzi, and Cristofolini 2016)) and calculated components of the iridial Lagrangian strain tensor at maximum pupillary constriction across the iris (Figure 2A). We observed strain patterns similar to that in an annular disc under axisymmetric radial contraction, with the *ϵ*_*xx*_ strain component distributed symmetrically about the nasal-temporal axis (*x*-axis) and *ϵ*_*yy*_ being symmetric about the superior-inferior axis (y-axis). The in-plane shear strain (*ϵ*_*xy*_) demonstrated an antisymmetric distribution across both x and y axes, with a 45° inclination (Figure 2A).

**Figure 2:**
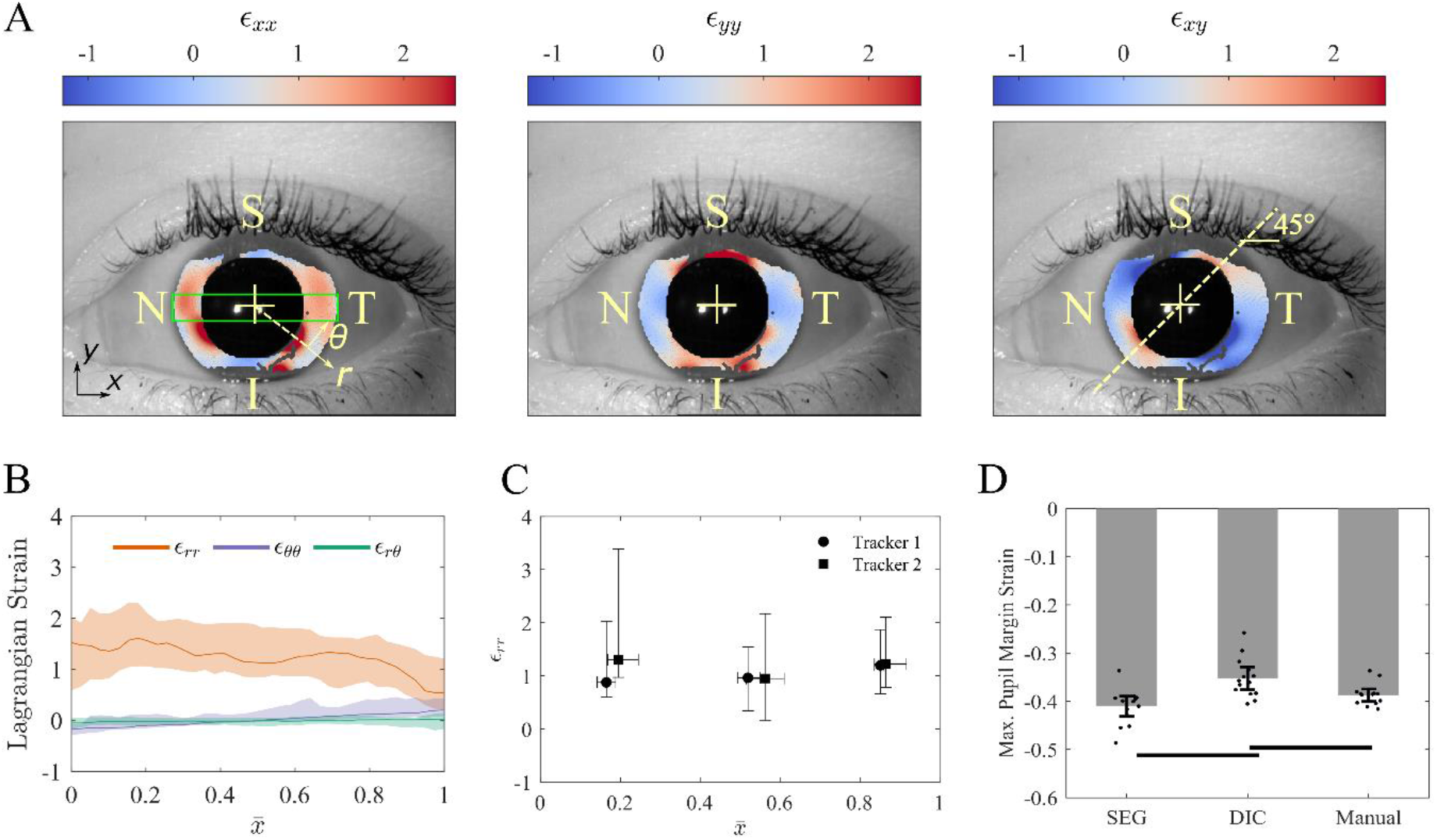
**(A)** Representative in-plane iridial Lagrangian strain field determined using digital image correlation (DIC). The colors indicate the strain at maximum pupillary constriction in the reference configuration. The strain fields demonstrate a symmetrical deformation, i.e. ***ϵ***_***xx***_ is essentially symmetric about the x-axis, ***ϵ***_***yy***_ is symmetric about the y-axis, and ***ϵ***_***xy***_ is diagonally antisymmetric. The color bar spans 95% of the CI of the data. S, N, I, T denote superior, nasal, inferior and temporal, respectively. **(B)** The spatial distribution of in-plane iridial strain components in a normalized coordinate system. The median and interquartile range (IQR; shaded areas) are shown for the ROI (green box in panel A left, with height equal to one-half of the pupil radius during the acclimation phase, and width equal to the limbus diameter), where 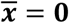 for the pupillary margin, and 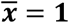 for the limbus. It is evident that there are significant deformations over the entire iris; for example, ***ϵ***_***rr***_ is 1.53 [0.59, 2.01] (median and IQR) at the pupillary margin. The median value of ***ϵ***_***rr***_ is essentially constant across much of the iris and then decrease to 0.54 [021, 1.22] at the limbus. As expected, ***ϵ***_***θθ***_ and ***ϵ***_***rθ***_ were small compared to ***ϵ***_***rr***_. ***ϵ***_***rθ***_ was negative at the pupillary margin (indicating sphincter constriction), and due to the symmetry of deformation, ***ϵ***_***rθ***_ was essentially zero. **(C)** We validated the DIC results by having two trackers annotate structural features manually to calculate ***ϵ***_***rr***_. By comparing the medians and IQR of ***ϵ***_***rr***_, it is evident that both trackers acquired similar results compared to DIC. In addition, the results of the two trackers were not different from each other. The vertical and horizontal error bars indicate IQR. **(D)** To further validate the DIC results, we measured the pupil strain by calculating the average pupil margin strain at maximum constriction (Manual) and compared it to pupil margin strain results from DIC analyses (DIC) and Daugman’s method (SEG). Results obtained by the three methods showed reasonable agreement, with the maximum difference of ~15% occurring between DIC and SEG (p<0.01). Error bars indicate 95% CI. The horizontal bars indicate p<0.05/3.

We selected a rectangular region of interest (ROI; green box in Figure 2A) centered on the pupil center to calculate the strain field for a standardized iris geometry in polar coordinates for all strain components. We calculated the median of each strain component in the ROI as a function of radial distance from the pupillary margin (see supplementary Figure S1), where *ϵ*_*xx*_ is essentially equivalent to *ϵ*_*rr*_, *ϵ*_*yy*_ to *ϵ*_*θθ*_, and *ϵ*_*xy*_ to *ϵ*_*rθ*_ (Figure 2A). Plotting these strain components vs. normalized distance (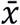) from the pupillary edge, we found that *ϵ*_*rr*_ was 1.53 [0.59, 2.01] (median and [interquartile range (IQR)]) at the pupillary edge and was almost uniform across the iris, with a localized decline to 0.54 [0.21, 1.22] close to the limbus (Figure 2B). In addition, both *ϵ*_*θθ*_ and *ϵ*_*rθ*_ were small compared to *ϵ*_*rr*_. *ϵ*_*θθ*_ was negative at the pupillary margin, consistent with sphincter constriction, and as expected due to symmetry in the iris deformation, *ϵ*_*rθ*_ was almost zero (Figure 2B).

To validate the DIC results, we compared them to strains obtained from two manual annotations, one for the spatial distribution of radial strain and the other for pupillary margin strain. First, two independent annotators manually tracked iridial features along the nasal-temporal axis (N-T; Figure 2A), from which we calculated the radial Lagrangian strain, *ϵ*_*rr*_, at maximum pupil constriction. The manual tracking results agreed with the DIC results, as demonstrated by comparing the median and IQR of the strains (Figure 2B & C). Further, the results of manual feature tracking were similar between the annotators (p=0.988; two-way ANOVA), and the strain values were not different between iris regions (p=0.407; two-way ANOVA).

We next manually calculated the maximum pupillary margin strain based on the change in the average diameter of the pupil during maximum constriction, calculated by averaging the diameter of the pupil along the nasal-temporal and superior-inferior axes (see supplementary Figure S2B). We compared these results to pupillary margin strain measured from DIC and segmentation/Daugman’s method (described above). The values of pupillary margin strain were generally consistent across the methods (Figure 2D), albeit with slightly different quantitative results (p=0.0001, two-way ANOVA), which was not dependent on the scanned eye (p=0.0817, two-way ANOVA). Only the strains from DIC showed a slight, albeit statistically significant, difference from the segmentation-based strains (*Δϵ*_*p*_ = 16.4%; p = 0.0017) and manual pupillary strains (*Δϵ*_*p*_ = 9.0%; p = 0.0021), while the segmentation-based vs. manual-based strain difference was not significant (*Δϵ*_*p*_ = 5.9%; p=0.025, i.e. greater than the Bonferroni-corrected significance level of 0.05/3).

### *In vivo* assessment of sphincter muscle traction

Next, we used experimentally-measured pupillary margin strains to evaluate iridial biomechanical properties *in vivo*. We modeled the iris using an eight-fold symmetric finite element mesh, with the inner pupillary elements representing the sphincter muscle (sphincter width *a*_*s*_ = 1mm; Figure 3 A&B) (Kaser-Eichberger et al. 2015; Moazed 2020). We performed multi-start data-fitting (Safa et al. 2021), using the measured mean maximum pupil margin strain as the target value and the model parameters being stromal modulus *E* (kPa), Poisson’s ratio *v*, and sphincter muscle traction *T*_*s*_ (kPa). Here, muscle traction was defined as the muscle contractile force divided by muscle cross-sectional area (normal to the pupil periphery). Interestingly, it was evident that the model fits were not sensitive to *v*, and that there was a linear correlation between *E* and *T*_*s*_, with *T*_*s*_: *E* ≈ 5 (Figure 3D). The *T*_*s*_: *E* ratio is important as it provides a basis for objective assessment of iris biomechanics from pupillary size changes, as discussed below.

**Figure 3:**
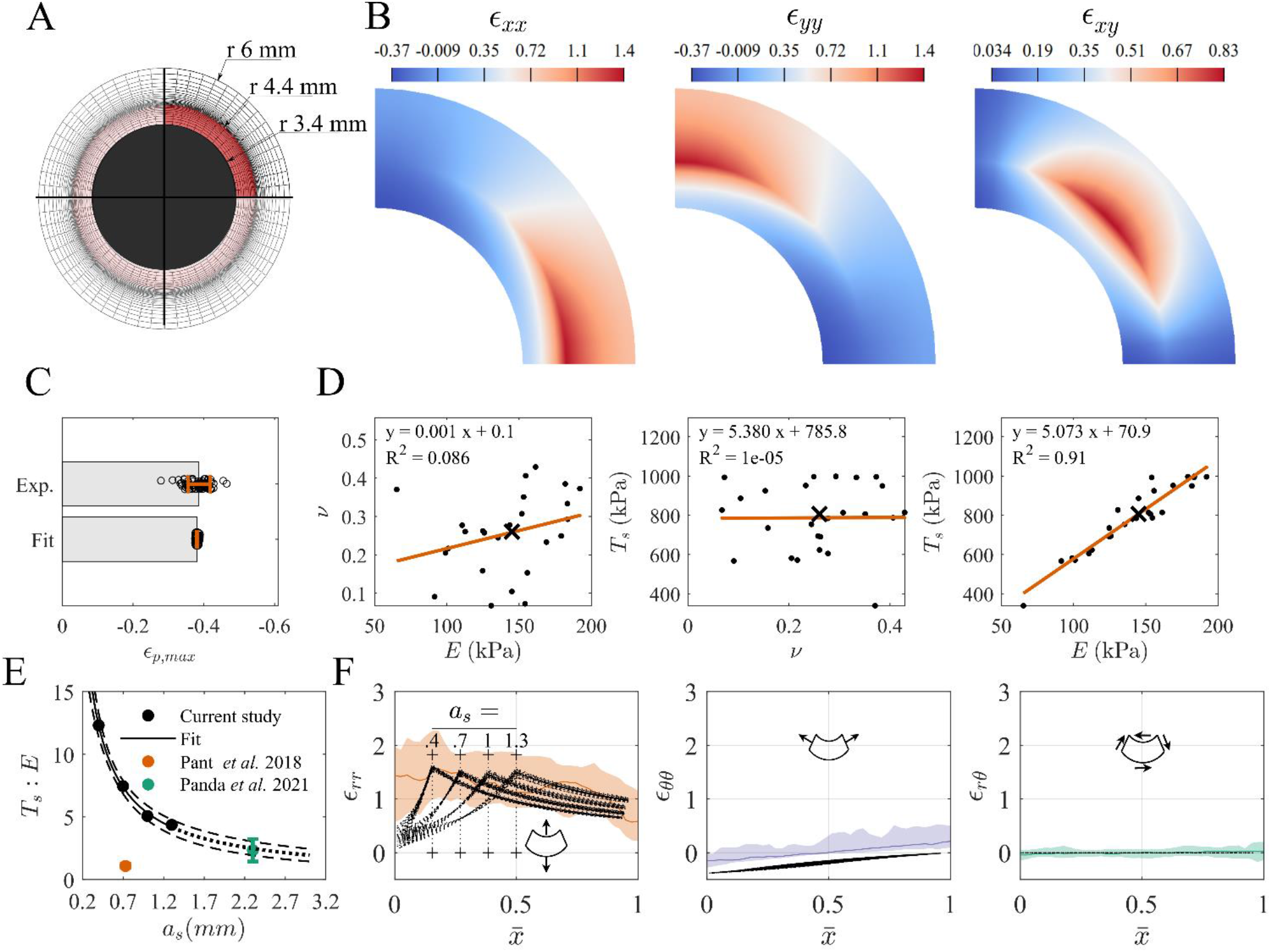
**(A)** The finite element [FE] mesh (sphincter width ***a***_***s***_ = **1 mm**) used to model iridial biomechanics. The active sphincter muscle is highlighted in red. **(B)** An example of the strain field obtained from FE simulations, where we leveraged the symmetry of the iris for numerical efficiency to model only a one-eighth wedge of the entire iris, i.e. half the iris thickness and a ¼ sector. **(C)** The model fitted the median experimental maximum pupillary strain very well (***a***_***s***_ = **1 mm**). Error bars are mean ± std deviation. **(D)** Cross-plots between the fitted model parameters, ***E, v***, and ***T***_***s***_, (***a***_***s***_ = **1 mm**, individual fit parameter values as solid dots, and the median as ‘×’) showing that the model fitting was insensitive to a change in ***v***, and interestingly that there was a strong linear correlation between ***E*** and ***T***_***s***_, with the latter being approximately five times the former. The linear regression results are shown in each panel, where ‘y’ corresponds to the parameter in the vertical axis and ‘x’ to the horizontal axis. **(E)** Repeating the data-fitting using different ***a***_***s***_ showed a nonlinear inverse correlation between the ***T***_***s***_: ***E*** ratio and ***a***_***s***_. The dashed-line indicates the 95% CI of the nonlinear regression analysis. The dotted-line indicates extrapolation of the model. The colored data points indicate values from the literature shown as mean ± std deviation. **(F)** Spatial distributions of Lagrangian strains (in polar coordinates) vs. normalized position across the iris, 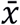. Symbols show results from the FE models for different ***a***_***s***_; colored lines and shaded regions show DIC measurements and 95% CI (repeated from Figure 2B). The model predictions and the experiments were in general agreement, especially when comparing the peak radial strains to the median experimental strains in the body of the iris; however, the shape of the radial strain was sensitive to ***a***_***s***_, where the peak radial strain occurred at the edge of the sphincter muscle (identified by vertical dotted lines and ‘+’ signs).

The force exerted by a muscle is dependent on its dimensions (Herlihy and Murphy 1973); therefore, we also conducted the above data-fitting while varying sphincter muscle width over a physiological range *a*_*s*_ = [0.4mm, 0.7mm, 1.3mm]. We observed that increasing *a*_*s*_ caused a decrease in the traction (*T*_*s*_: *E* ratio) needed to achieve the same pupillary strain, from 13 to 4. The relation between *T*_*s*_: *E* and *a*_*s*_ was non-linear and could be fit by the following empirical relation:

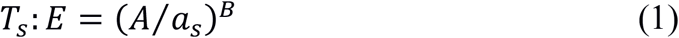

where *T*_*s*_: *E* is non-dimensional, *a*_*s*_ is in mm, A = 6.197 [3.723, 8.671] mm (mean [95% confidence interval], and B = 0.916 [0.768, 1.064] is non-dimensional (Figure 3E).

A further effect of changing *a*_*s*_ was alteration in the spatial distribution of iridial strains. The simulated strain responses for different *a*_*s*_ values were compared to each other and to the values experimentally measured using DIC, as described above (Figure 2B). We observed that changing *a*_*s*_ affected the spatial distribution of the radial strain (*ϵ*_*rr*_); however, it did not change the spatial distribution of *ϵ*_*θθ*_ and *ϵ*_*rθ*_ (Figure 3F), despite the simulations having different sets of material parameter values. Further, the peak *ϵ*_*rr*_ was the same for all the models and agreed with the DIC results. However, within the sphincter muscle, *ϵ*_*rr*_ had a different distribution compared to the experimentally-measured strains, with the model having a smaller *ϵ*_*rr*_ value compared to the experimental data. There was agreement between the experimental and modelled *ϵ*_*θθ*_ and *ϵ*_*rθ*_, albeit with a negative shift in *ϵ*_*θθ*_ of the models compared to the experimental values. Specifically, the model’s predicted *ϵ*_*θθ*_ was c. −40%, which is consistent with the strain determined from changes in pupillary diameter, suggesting that the DIC underestimated *ϵ*_*θθ*_.

## Discussion

The iris plays a central role in primary angle-closure glaucoma (PACG). Worldwide, PACG is the second most prevalent form of glaucoma, although in some regions, primarily in parts of East Asia, PACG is the most prevalent form (Cheng et al. 2014). Current risk assessment in PACG patients is based on precise anatomical measurements of the anterior chamber and iris, usually by optical coherence tomographic (OCT) imaging (Friedman et al. 2012; You et al. 2021), yet the predictive power of such techniques is poor (He et al. 2019). This motivates the development of novel techniques for identification and assessment of PACG risk factors.

Iris biomechanical properties have been largely ignored as potential risk factors for PACG despite their likely importance. There are two challenges in incorporating iris biomechanics into clinical management of PACG. First, knowledge about iris biomechanical properties is scarce. Second, there are currently no clinical techniques to measure iris biomechanics. Ideally such techniques would be inexpensive, i.e. suitable for patients in less economically-developed settings. Here we repurposed a publicly available biometrics data set to track the deformations of the iris during the pupillary light reflex (PLR) and hence analyze the *in vivo* biomechanical properties of the human iris. Inducing and measuring PLR is accessible and reproducible, and thus our approach is amenable to future translational studies of iris biomechanics.

Our data showed that the sphincter muscle traction (*T*_*s*_) and iris stroma’s stiffness (*E*) are linearly correlated, with a mean *T*_*s*_: *E* ratio of 5 (range 4-13, depending on sphincter muscle width). The correlation between *T*_*s*_ and *E* is consistent with previous studies, where in non-glaucomatous human subjects, Pant *et al.* (Pant et al. 2018) estimated *T*_*s*_: *E* = 1.08 ± 0.16, and Panda *et al.* estimated *T*_*s*_: *E* = 2.34 ± 0.90 (Panda et al. 2021) (Figure 3E), while in subjects with a history of PACG, the sphincter muscle was determined to be relatively weaker, with *T*_*s*_: *E* = 0.38 ± 0.10 (Pant et al. 2018) and *T*_*s*_: *E* = 1.63 ± 0.56 (Panda et al. 2021).

It is of interest to note that the *T*_*s*_: *E* values from previous studies are notably smaller than our estimate of *T*_*s*_: *E* (Figure 3E). There are multiple interrelated factors that likely influence this difference, as follows.

- *Sphincter muscle size*. Panda *et al.* used a sphincter muscle size much larger than ours. Their muscle size was measured in porcine eyes, yet there are notable anatomic differences between pig and human irides (e.g. elliptical pupils), suggesting an over-estimation of human muscle size in their study. Interestingly, our empirical relation (Equation 1) is consistent with the *T*_*s*_: *E* value that they report (Figure 3E), i.e. if Panda *et al.* had used smaller sphincter width in their study, they may have had arrived at a similar *T*_*s*_: *E* ratio as us. Unfortunately, there are currently no *in vivo* methods for determining sphincter muscle dimensions, motivating the development of techniques for assessing sphincter muscle size in human subjects.
- *Methodological differences*: Pant *et al.* found a *T*_*s*_: *E* ratio 3-fold smaller than ours, even though the sphincter muscle width that they used (~0.73 mm) lay within our range (*a*_*s*_ = 0.4-1.3mm). However, they used less extreme lighting conditions to induce PLR, resulting in smaller iridial radial strains than we observed (~12% vs. 100%). Presumably, this means that the sphincter muscle was not maximally stimulated in their study, emphasizing the importance of methodological details.
- *Genetic background:* Pant *et al.* studied an Indian population, and Panda *et al.* studied a Singaporean one (Indian/Chinese ethnicity), while our data originated in Poland. Although patient demographics were not available for our population, it is highly likely that subjects were of European descent. We speculate that populations of European descent have a larger *T*_*s*_: *E* than Asian populations, due to several related observations. First, PACG is more prevalent in Asia, including both India and Singapore, than in the rest of the world (Cheng et al. 2014). Second, PACG patients have stiffer irides (Pant et al. 2018; Panda et al. 2021). Clearly further study is required to evaluate whether there are differences in iridial biomechanics between different populations. Identification of such differences would complement established anatomical risk factors in genetically diverse clinical populations.

We showed that the iris experiences radial strains (*ϵ*_*rr*_) of greater than 100% during PLR (Figure 2), which is consistent with previous reports using manual feature tracking (Pamplona, Oliveira, and Baranoski 2009; Wyatt 2000). Further, we observed that the circumferential strain (*ϵ*_*θθ*_) was also significant (Figure 2A and B). At the pupil edge, *ϵ*_*θθ*_ calculated from the FE model matched *ϵ*_*θθ*_ computed from tracking pupil diameter experimentally, but not values of *ϵ*_*θθ*_ measured by DIC (Figure 3F). Likely this discrepancy is due to the radial orientation of iris surface features and the large deformations in the radial direction, complicating DIC imaging. Higher resolution imaging would be needed to more accurately calculate the circumferential strains of the iris by DIC.

This study was subject to several limitations. For example, we did not include the dilator muscle in our analyses. However, the effects of dilator muscle traction during PLR are minimal (Loewenfeld and Lowenstein 1999). Additionally, since we used 2D images, there were potential confounding factors due to the curvature of the iris, distortions due to corneal refraction, and reflected light on the cornea. For example, we observed a subtle decline in the median radial strain near the limbus (Figure 2B), potentially due distortion due to corneal refraction in this region. In addition, corneal reflections added noise which complicated feature tracking and DIC (see supplementary Figure S4). Future studies could benefit from using 3D imaging modalities (e.g., OCT and elastography) to validate deformations measured from NiR imaging. 3D imaging of the iris could also be helpful to identify additional *in vivo* mechanical properties, such as anisotropic material properties and Poisson’s ratio, which our model could not evaluate (Panda et al. 2021).

In conclusion, we measured iridial deformations and determined tissue mechanical properties *in vivo* using imaging of the pupillary light reflex (PLR) and finite element modeling. Our technique for measuring iris biomechanics is simple and does not require specialized devices, and therefore has significant potential for clinical translation. This study establishes proof-of-concept for using pupillography during the PLR to functionally assess iris biomechanics *in vivo*, of interest in evaluating iris biomechanics’ role in glaucoma.

## Materials and Methods

### Pupillary light reflex data set

To assess the tissue deformations induced by the activation of the iris sphincter muscle, we used the publicly available *Warsaw-BioBase-Pupil-Dynamics v3* data set (Kinnison et al. 2019), which includes 163 videos (each 30 seconds long, acquired at 25 Hz) of PLR from 42 subjects of ages 20-50 years. Each eye scan video has a unique code; e.g. *10066left2* denotes the second scan of subject 10066’s left eye. To obtain scans, the subject’s head was placed in a large shaded box to prevent penetration of ambient light, and built-in LEDs were used to induce the PLR. Images were acquired in the near-infrared (NiR) using a custom system (IrisCube (Czajka 2015)). NiR imaging is standard practice in pupilography (Kelbsch et al. 2019), where light with a wavelength less than 800 nm is absent, allowing imaging in darkness and detection of pupillary reflexes independent of the stimulus lighting. Both left (OS) and right (OD) eyes of subjects were scanned twice. Scans included a 15-second acclimation phase in the dark (dark adaptation), a 5 second exposure of the eye to LED light, followed by 10 seconds of darkness (Figure 1A). The 5 second exposure to visible light induced pupillary constriction and the elimination of this stimulus allowed for partial pupil recovery. We note that full pupil size recovery can be achieved with a longer period of darkness after the light stimulus (Ba-Ali et al. 2020; Joyce et al. 2018); fortunately, the lack of full recovery in this data set did not affect our analysis, since we were only interested in maximum pupil constriction.

### Pupil and limbus segmentation and deformation

To assess the deformation of the pupil and limbus, we used an automated algorithm. Given the enormous volume of data, we analyzed every tenth image in the videos, resulting in an effective 2.5 Hz frequency, equivalent to 400 msec temporal resolution. Unless otherwise specified, all analyses were performed in Matlab. Specifically, to measure pupil edge diameter, we used a custom pixel intensity-based threshold segmentation of the pupil, where we first applied a median filter (*medfilt2()* function; window size = [3 pixel × 3 pixel]) to reduce image noise, followed with a binarization function based on Otsu’s method (*imbinarize()* function) with a 0.1 threshold. Next, to obtain a final pupil mask, we performed an erosion and dilation routine (*imerode()* and *imdilate()* functions) with a 2 pixels-wide square morphological element (*strel()* function), and fill hole (*imfill()* function). We then calculated the average pupil radius as 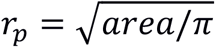. To measure limbus radius, we used a publicly available implementation of Daugman’s method in Matlab (Sivaraman 2021; Daugman 2004). The outputs of this step were the fitted radii of the pupil and limbus. We calculated the ratio of the pupillary (*p*) to limbal (*l*) radii as:

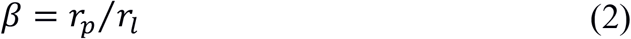

We also calculated the Lagrangian strain of both the pupillary margin and limbus as:

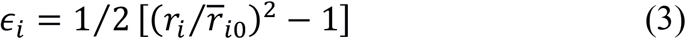

where 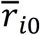 is the average value of *r*_*i*_ over the initial 15-second acclimation phase (Figure 1C), and *i* = *p* for pupil and *i* = *l* for limbus.

We calculated the maximum pupillary margin strain, used in the finite element analysis, as the mean strain over the interval 17-20 sec. We averaged the test-retest scans for each eye, and then calculated the mean, standard deviation, and 95% confidence interval of the maximum pupillary margin strain for the entire data set. As quality control, we identified failed segmentations by performing a *post hoc* outlier identification, where the segmentations having maximum pupillary margin strain values more than three times the standard deviation away from the mean were excluded from the analysis. To test whether repeated scans of each eye or the eye’s anatomical placement (OS/OD) affected the PLR, we used a linear mixed-effects model (LME) with the pupillary margin strain as the observed parameter, fixed effects being the order of scan (scan 1 and scan 2) and anatomical placement (OS and OD), and random effect being the anatomical placement grouped based on subject (significance level *α* = 0.05).

### Spatial distribution of strain and digital image correlation

We calculated the deformation in the iris stroma during PLR using digital image correlation (DIC). We conducted the DIC analysis using Vic-2D software (Correlated Solution, Irmo, SC, USA) on n = 17 videos from nine unique eyes from seven subjects. Some of the analyzed videos were repeat scans of the same eye; however, due to the randomness of the gaze, blinking, and corneal reflection patterns, we treated each video as an independent sample for the DIC analysis.

The images (768 pixels wide × 576 pixels high) were loaded into Vic-2D using the tagged image file format (tiff). We conducted an incremental correlation (subset size of 31 pixels and step size of 4) on a manually traced reference ROI around the iris that excluded the pupil and eyelids from the analysis. The matchability threshold was set at 0.1 pixels, and the Lagrangian strain was calculated in a post-processing step with a filter size of 15 pixels. We analyzed the images after the beginning of light stimulation, i.e., during pupillary constriction. To avoid the effect of blinking, which could terminate the DIC tracking, we manually excluded images in which blinking occurred. For consistency, we used the same protocol for all DIC analyses.

We evaluated the spatial distribution of the Lagrangian normal and shear strains along the nasal-temporal axis at maximum pupillary constriction. The strain fields near the superior and inferior regions were not reliable due to coverage by the eyelids (see supplementary Figure S3). We evaluated the strain along the nasal-temporal axis by calculating the median of the strains along the superior-inferior axis in a rectangular ROI that passed through the pupillary center, of height one-quarter of the pupillary diameter and width equal to the limbus diameter (green box in Figure 3A and supplementary Figure S1). To maintain a consistent coordinate system for all the strain fields, we used a normalized distance from the pupil margin in which the pupillary margin had a coordinate value of zero 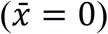, and the limbus had a value of 1 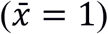. We conducted a *post hoc* outlier identification analysis based on the Hausdorff distance (Danziger 2021) of the strain component curve vs. 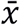 and excluded four videos from the DIC analysis; however, this had a minimal effect on the results (see supplementary Figure S5).

To validate the DIC results, we used two procedures. First, two separate annotators (trackers) manually tracked eight points along the nasal-temporal axis using ImageJ (Schneider, Rasband, and Eliceiri 2012) in a subset of the videos analyzed by DIC (n=8). We selected approximately equally distanced points along the nasal-temporal axis to divide the area between limbus and pupil into three roughly equal parts (see supplementary Figure S2). However, the choice of points was limited by the traceability of features with unaided human vision. We calculated the Lagrangian strain along the nasal-temporal axis at maximum pupillary constriction and conducted a two-way ANOVA, where the factors were trackers and regions (α=0.05). Second, we compared the pupillary margin strains measured from segmentation (*ϵ*_*p,max*_ SEG) to the pupillary margin strain measured using a virtual tensometer in Vic-2D (*ϵ*_*p,max*_ DIC). For the latter comparison we also added another set of manual measurements of pupillary strain at maximum constriction (*ϵ*_*p,max*_ Manual) (n=17), where we calculated the average of the Lagrangian pupillary margin strain at three time-points (frames 425/750, 463/750, 500/750) according to Equation 3, with the onset of light stimulation (frame 375/750) being the reference (see supplementary Figure S2). Next, we conducted a two-way ANOVA with the factors being analysis method (SEG, DIC, and Manual) and the identification code of each eye scan (α=0.05), followed by a *post hoc* paired t-test with Bonferroni correction (α=0.05/3). The pupillary margin strain could not be calculated for three movies because the DIC algorithm failed to pass internal quality control thresholds at the pupillary margin; therefore, we excluded those samples from the ANOVA.

### Finite element modeling of the iris

We created an idealized semi-2D model of the iris, composed of an eight-fold symmetric portion of a disc under plane-stress boundary conditions, motivated by the assumption that anterior and posterior chambers were at the same pressure, resulting in zero net force loading. Details of the model boundary conditions are shown in supplementary Figure S6. We took the iris during the acclimation phase (Figure 1D) as the reference state. The outer radius of the model was 6 mm (Bergmanson and Martinez 2017), and the thickness of the model was 0.17 mm (based on average iris thickness of 0.34 mm (Marchini et al. 1986)). We set the inner radius of the model (pupillary radius) to 3.4 mm, which was calculated based on the outer radius of the iris and the average ratio of the pupillary and limbal radii during the acclimation period (*β*, Figure 1C). We used 2250 hexahedral elements (HEX8) to generate the mesh, based on a preliminary mesh density sensitivity analysis.

We modeled the iris’ mechanical response using a hyperelastic stromal substance with embedded uniaxial active traction elements to represent the sphincter muscle (Figure 3A). To simplify the model, we assumed that the sphincter was distributed across the radius in a ring of thickness *a*_*s*_ = 0.4 – 1.3 mm (Kaser-Eichberger et al. 2015; Moazed 2020) and modeled its traction as a 1D active material along the periphery of the pupil edge, i.e., the Cauchy stress due to the sphincter muscle was (Pant et al. 2018):

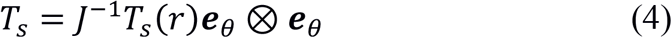

Here, *T*_*s*_ is the magnitude of the sphincter muscle traction, *r* is the distance from the pupil center, ***e***_*θ*_ is the circumferential unit vector along the sphincter muscle in the deformed state, and *J* is the Jacobian of the deformation gradient tensor.

Further, we described the mechanical response of the stroma using a compressible neo-Hookean constitutive relation:

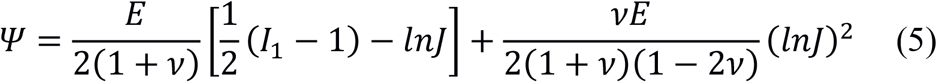

where *Ψ* is Helmholtz’s free energy, *I*_1_ is the first invariant of the Cauchy-Green deformation tensor; *E* is Young’s modulus (stiffness), and *v* is Poisson’s ratio. The model was implemented and solved using the FEBio software suite (FEBio v 3.1 (Maas et al. 2012)), and an example of the finite element model’s output is shown in Figure 3B.

### Parameter identification

We used the absolute value of the difference between the experimental and the modeled pupillary margin strain at maximum constriction (taken as the average over the period 17-20 sec) to perform data-fitting, with parameters *E*, *v* and *T*_*s*_. We used a multi-start optimization method (Safa et al. 2021) with a grid size of 25, to eliminate bias to one initial guess. We set a wide search space with 0 < *E* < 1000 kPa, 0 < *v* < 0.5, and 0 < *T*_*s*_ < 1000 kPa, which was informed by values previously reported in the literature (Panda et al. 2021; Pant et al. 2018; Whitcomb et al. 2011; 2009; Ye et al. 2021). We performed the data-fitting based on a baseline sphincter muscle width of 1 mm, and due to the sensitivity of the results to the assumed value of sphincter muscle width, we repeated the simulations for *a*_*s*_ = [0.4 mm, 0.7 mm, 1.0 mm, 1.3 mm]. Using the resulting fitted values, we nonlinearly regressed the ratio of sphincter traction to stroma stiffness vs. *a*_*s*_.

## Acknowledgments

We acknowledge our funding sources NIH-NEI (R01 EY031710, CRE), the Georgia Research Alliance (CRE), and the BrightFocus Foundation (postdoctoral fellowship G2021005F, BNS).

## References

Amini, Rouzbeh, Julie E Whitcomb, Muhammad K Al-Qaisi, Taner Akkin, Sara Jouzdani, Syril Dorairaj, Tiago Prata, et al. 2012. “The Posterior Location of the Dilator Muscle Induces Anterior Iris Bowing during Dilation, Even in the Absence of Pupillary Block.” Investigative Ophthalmology & Visual Science 53 (3): 1188–94. https://doi.org/10.1167/iovs.11-8408.

Ba-Ali, Shakoor, Adam Elias Brøndsted, Henrik Ullits Andersen, Poul Jennum, and Henrik Lund-Andersen. 2020. “Pupillary Light Responses in Type 1 and Type 2 Diabetics with and without Retinopathy.” Acta Ophthalmologica 98 (5): 477–84. https://doi.org/10.1111/aos.14348.

Bergmanson, Jan PG, and Jesus G. Martinez. 2017. “Size Does Matter: What Is the Corneo-Limbal Diameter?” Clinical and Experimental Optometry 100 (5): 522–28. https://doi.org/10.1111/cxo.12583.

Chang, Dolly S., Li Xu, Michael V. Boland, and David S. Friedman. 2013. “Accuracy of Pupil Assessment for the Detection of Glaucoma.” Ophthalmology 120 (11). https://doi.org/10.1016/j.ophtha.2013.04.012.

Cheng, Jin-Wei, Ying Zong, You-Yan Zeng, and Rui-Li Wei. 2014. “The Prevalence of Primary Angle Closure Glaucoma in Adult Asians: A Systematic Review and Meta-Analysis.” PLOS ONE 9 (7): e103222. https://doi.org/10.1371/journal.pone.0103222.

Czajka, A. 2015. “Pupil Dynamics for Iris Liveness Detection.” IEEE Transactions on Information Forensics and Security 10 (4): 726–35. https://doi.org/10.1109/TIFS.2015.2398815.

Danziger, Zachary. 2021. “Hausdorff Distance.” 2021. https://www.mathworks.com/matlabcentral/fileexchange/26738-hausdorff-distance.

Daugman, J. 2004. “How Iris Recognition Works.” IEEE Transactions on Circuits and Systems for Video Technology 14 (1): 21–30. https://doi.org/10.1109/TCSVT.2003.818350.

Friedman, David S., Paul J. Foster, Tin Aung, and Mingguang He. 2012. “Angle Closure and Angle-Closure Glaucoma: What We Are Doing Now and What We Will Be Doing in the Future.” Clinical & Experimental Ophthalmology 40 (4): 381–87. https://doi.org/10.1111/j.1442-9071.2012.02774.x.

He, Mingguang, Yuzhen Jiang, Shengsong Huang, Dolly S. Chang, Beatriz Munoz, Tin Aung, Paul J. Foster, and David S. Friedman. 2019. “Laser Peripheral Iridotomy for the Prevention of Angle Closure: A Single-Centre, Randomised Controlled Trial.” Lancet (London, England) 393 (10181): 1609–18. https://doi.org/10.1016/S0140-6736(18)32607-2.

Herlihy, Jeremiah T., and Richard A. Murphy. 1973. “Length-Tension Relationship of Smooth Muscle of the Hog Carotid Artery.” Circulation Research 33 (3): 275–83. https://doi.org/10.1161/01.RES.33.3.275.

Joyce, Daniel S., Beatrix Feigl, Graham Kerr, Luisa Roeder, and Andrew J. Zele. 2018. “Melanopsin-Mediated Pupil Function Is Impaired in Parkinson’s Disease.” Scientific Reports 8 (1): 7796. https://doi.org/10.1038/s41598-018-26078-0.

Kaser-Eichberger, Alexandra, Falk Schrödl, Andrea Trost, Clemens Strohmaier, Barbara Bogner, Christian Runge, Karolina Motloch, et al. 2015. “Topography of Lymphatic Markers in Human Iris and Ciliary Body.” Investigative Opthalmology & Visual Science 56 (8): 4943. https://doi.org/10.1167/iovs.15-16573.

Kelbsch, Carina, Torsten Strasser, Yanjun Chen, Beatrix Feigl, Paul D. Gamlin, Randy Kardon, Tobias Peters, et al. 2019. “Standards in Pupillography.” Frontiers in Neurology 10 (February): 129. https://doi.org/10.3389/fneur.2019.00129.

Kinnison, Jeffery, Mateusz Trokielewicz, Camila Carballo, Adam Czajka, and Walter Scheirer. 2019. “Learning-Free Iris Segmentation Revisited: A First Step Toward Fast Volumetric Operation Over Video Samples.” In 2019 International Conference on Biometrics (ICB), 1–8. https://doi.org/10.1109/ICB45273.2019.8987377.

Loewenfeld, Irene E., and Otto Lowenstein. 1999. The Pupil: Anatomy, Physiology, and Clinical Applications. Butterworth-Heinemann.

Maas, Steve A., Benjamin J. Ellis, Gerard A. Ateshian, and Jeffrey A. Weiss. 2012. “FEBio: Finite Elements for Biomechanics.” Journal of Biomechanical Engineering 134 (1): 011005–011005. https://doi.org/10.1115/1.4005694.

Marchini, M., M. Morocutti, A. Ruggeri, M. H.J. Koch, A. Bigi, and N. Roveri. 1986. “Differences in the Fibril Structure of Corneal and Tendon Collagen. An Electron Microscopy and x-Ray Diffraction Investigation.” Connective Tissue Research 15 (4): 269–81. https://doi.org/10.3109/03008208609001985.

Moazed, Kambiz Thomas. 2020. The Iris: Understanding the Essentials.

Newsome, David A., and Irene E. Loewenfeld. 1971. “Iris Mechanics II. Influence of Pupil Size on Details of Iris Structure.” American Journal of Ophthalmology 71 (2): 553–73. https://doi.org/10.1016/0002-9394(71)90133-4.

Omelina, Lubos, Jozef Goga, Jarmila Pavlovicova, Milos Oravec, and Bart Jansen. 2021. “A Survey of Iris Datasets.” Image and Vision Computing 108 (April): 104109. https://doi.org/10.1016/j.imavis.2021.104109.

Palanca, Marco, Gianluca Tozzi, and Luca Cristofolini. 2016. “The Use of Digital Image Correlation in the Biomechanical Area: A Review.” International Biomechanics 3 (1): 1–21. https://doi.org/10.1080/23335432.2015.1117395.

Pamplona, Vitor F., Manuel M. Oliveira, and Gladimir V. G. Baranoski. 2009. “Photorealistic Models for Pupil Light Reflex and Iridal Pattern Deformation.” ACM Transactions on Graphics 28 (4): 1–12. https://doi.org/10.1145/1559755.1559763.

Panda, Satish K., Royston K. Y. Tan, Tin A. Tun, Martin L. Buist, Monisha Nongpiur, Mani Baskaran, Tin Aung, and Michaël J. A. Girard. 2021. “Changes in Iris Stiffness and Permeability in Primary Angle Closure Glaucoma.” Investigative Ophthalmology & Visual Science 62 (13): 29. https://doi.org/10.1167/iovs.62.13.29.

Pant, Anup Dev, Priyanka Gogte, Vanita Pathak-Ray, Syril K. Dorairaj, and Rouzbeh Amini. 2018. “Increased Iris Stiffness in Patients With a History of Angle-Closure Glaucoma: An Image-Based Inverse Modeling Analysis.” Investigative Ophthalmology & Visual Science 59 (10): 4134–42. https://doi.org/10.1167/iovs.18-24327.

Quigley, H. A., D. M. Silver, R. J. Plyler, and D. S. Friedman. 2008. “The Iris Loses Half Its Volume During Pupil Dilation: A New Risk Factor for Angle Closure Glaucoma.” Investigative Ophthalmology & Visual Science 49 (13): 5100–5100.

Safa, B. N., M. H. Santare, C. R. Ethier, and D. M. Elliott. 2021. “Identifiability of Tissue Material Parameters from Uniaxial Tests Using Multi-Start Optimization.” Acta Biomaterialia, January. https://doi.org/10.1016/j.actbio.2021.01.006.

Schneider, Caroline A, Wayne S Rasband, and Kevin W Eliceiri. 2012. “NIH Image to ImageJ: 25 Years of Image Analysis.” Nature Methods 9 (7): 671–75. https://doi.org/10.1038/nmeth.2089.

Sivaraman, Anirudh. 2021. “Iris Segmentation Using Daugman’s Integrodifferential Operator.” 2021. https://www.mathworks.com/matlabcentral/fileexchange/15652-iris-segmentation-using-daugman-s-integrodifferential-operator.

Tan, Royston K. Y., Xiaofei Wang, Anita S. Y. Chan, Monisha Esther Nongpiur, Craig Boote, Shamira A. Perera, and Michaël J. A. Girard. 2019. “Permeability of the Porcine Iris Stroma.” Experimental Eye Research 181 (April): 190–96. https://doi.org/10.1016/j.exer.2019.02.005.

Whitcomb, Julie E, Rouzbeh Amini, Narendra K Simha, and Victor H Barocas. 2011. “Anterior-Posterior Asymmetry in Iris Mechanics Measured by Indentation.” Experimental Eye Research 93 (4): 475–81. https://doi.org/10.1016/j.exer.2011.06.009.

Whitcomb, Julie E, Vincent A Barnett, Timothy W Olsen, and Victor H Barocas. 2009. “Ex Vivo Porcine Iris Stiffening Due to Drug Stimulation.” Experimental Eye Research 89 (4): 456–61. https://doi.org/10.1016/j.exer.2009.04.014.

World Health Organization. 2019. World Report on Vision. World Health Organisation. Vol. 214. 14. https://www.who.int/publications/i/item/world-report-on-vision.

Wyatt, H J. 2000. “A ‘minimum-Wear-and-Tear’ Meshwork for the Iris.” Vision Research 40 (16): 2167–76. https://doi.org/10.1016/s0042-6989(00)00068-7.

Ye, Shuling, Yuheng Zhou, Chenhong Bao, Yulei Chen, Fan Lu, and Dexi Zhu. 2021. “In Vivo Non-Contact Measurement of Human Iris Elasticity by Optical Coherence Elastography.” Journal of Biophotonics, May, e202100116. https://doi.org/10.1002/jbio.202100116.

You, Shuqi, Zhiqiao Liang, Kangyi Yang, Yu Zhang, Julius Oatts, Ying Han, and Huijuan Wu. 2021. “Novel Discoveries of Anterior Segment Parameters in Fellow Eyes of Acute Primary Angle Closure and Chronic Primary Angle Closure Glaucoma.” Investigative Ophthalmology & Visual Science 62 (14): 6. https://doi.org/10.1167/iovs.62.14.6.

