## Supplementary document for "In vivo Biomechanical Assessment of Iridial Deformations and Muscle Contractions in Human Eyes"

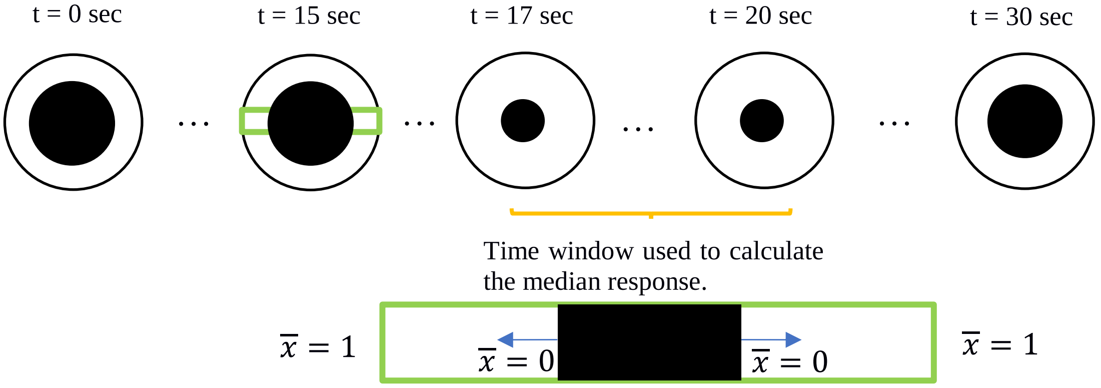


**Figure S1**: A schematic of the procedure for calculating the median strain in the region of interest (ROI, green box). To calculate the strain field at maximum pupillary constriction, we evaluated the median field over the time interval 17-20 sec (yellow bracket) of the DIC strain fields (i.e., $\overline{\boldsymbol{E}}(x, y) = \text{median}\boldsymbol{E}(x, y, t)$). We then calculated the spatial median of the strain component in a normalized coordinate system (i.e., $E(\bar{x}) = \text{median}\overline{\boldsymbol{E}}(x, y)$), where, as shown in the enlarged ROI green box in the bottom of the figure, $\overline{x} = 0$ corresponded to the location of the pupillary margin and $\overline{x} = 1$ corresponded to the limbal margin position on each side of the eye.

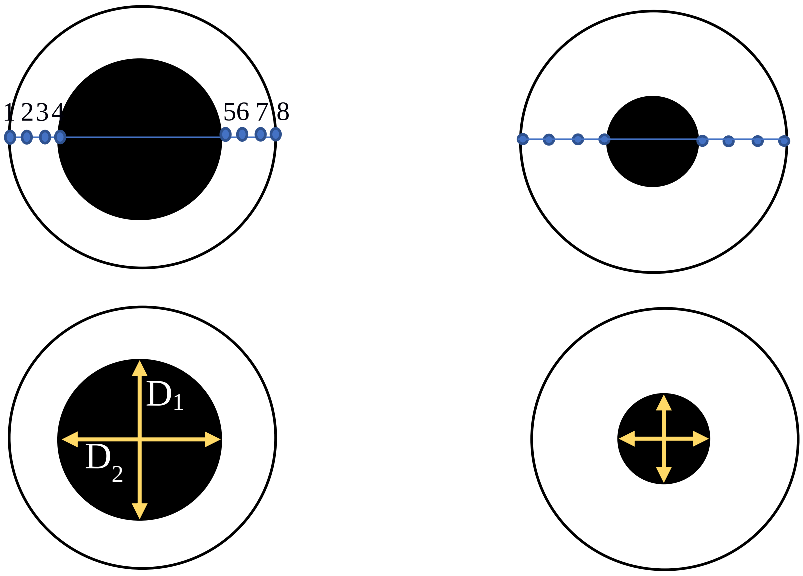


**Figure S2: (A)** Schematic of the procedure for manual evaluation of the radial strain using 8 points along the nasal-temporal axis of the eye. **(B)** Manual evaluation of the average pupil diameter where $D_{average} = (D_1 + D_2)/2$.

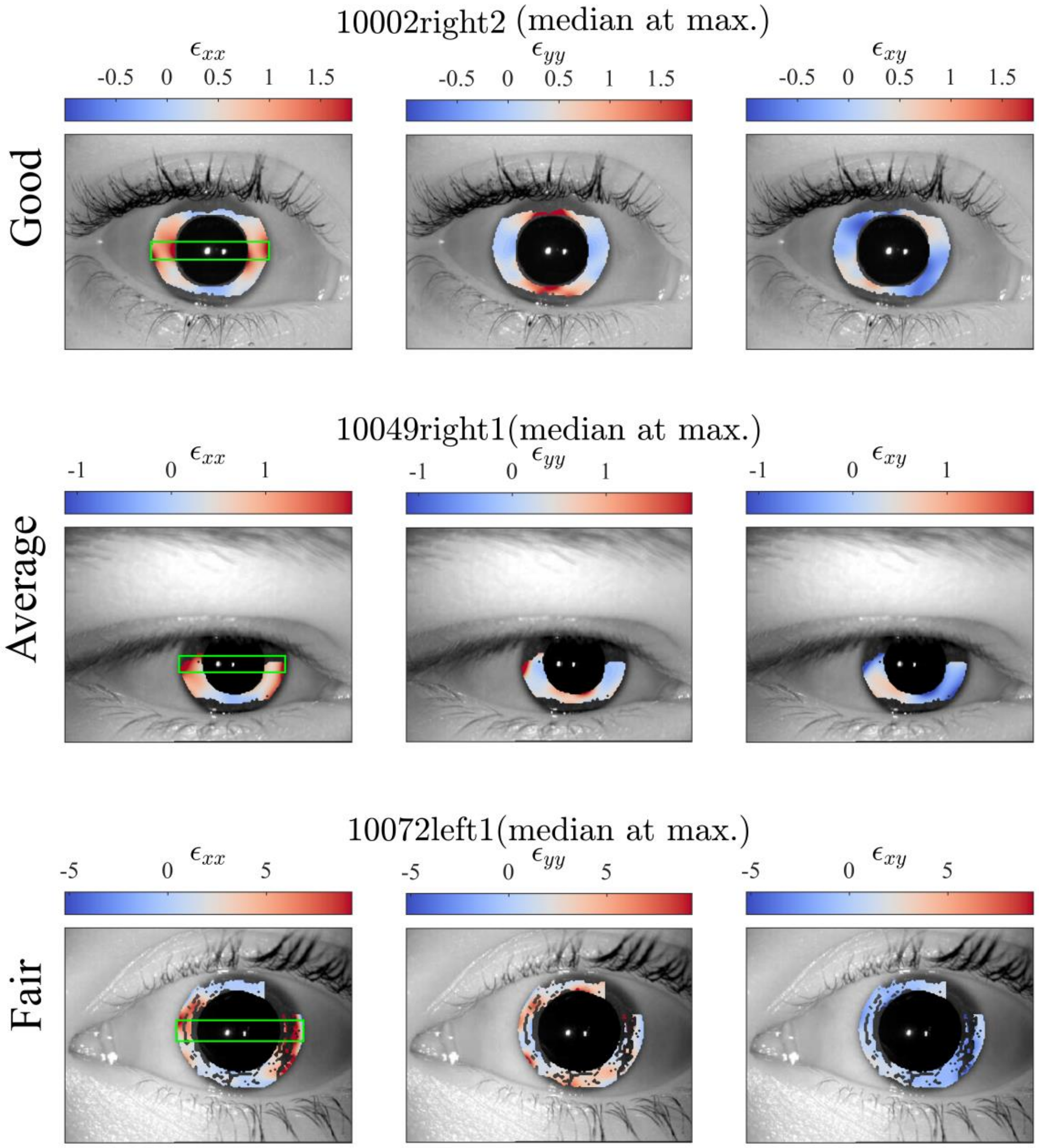


**Figure S3**: Three representative results of the DIC analysis demonstrating varying result quality. The DIC analysis's quality depends on the prominence of iris features, gaze direction, blinking, and corneal reflections. The plotted quantities are temporal median strain fields at maximum pupillary constriction ("Median at max."). Grey regions indicate locations where the DIC algorithm failed.

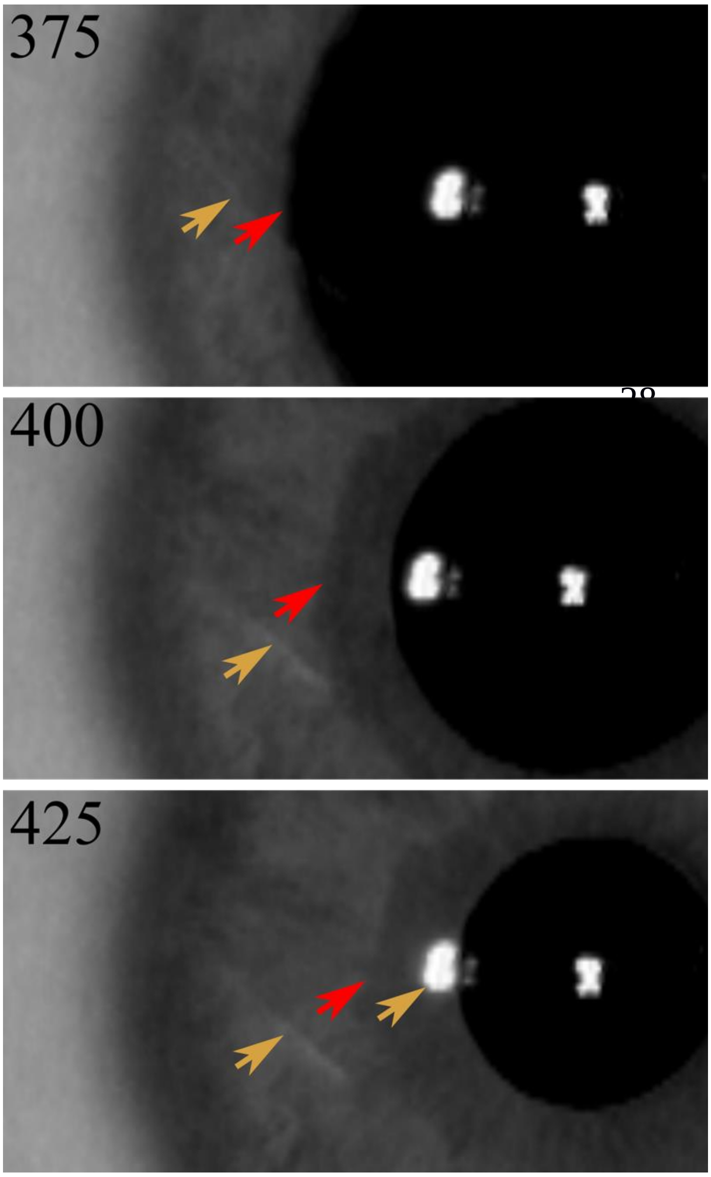


**Figure S4:** An example of a scan with imaging artifacts that impeded DIC analysis (10066left2). We have included three images from the same scan (raw image at top and zoomed-in views around the pupil at bottom) at the beginning of light exposure (frame 375), during constriction (frame 400), and during maximum constriction (frame 425). The red arrows indicate a region near the pupillary edge which appears to be unfolding from beneath the iris, implying likely 3D deformation. The orange arrows indicate corneal reflections that hinder convergence of the DIC algorithm.

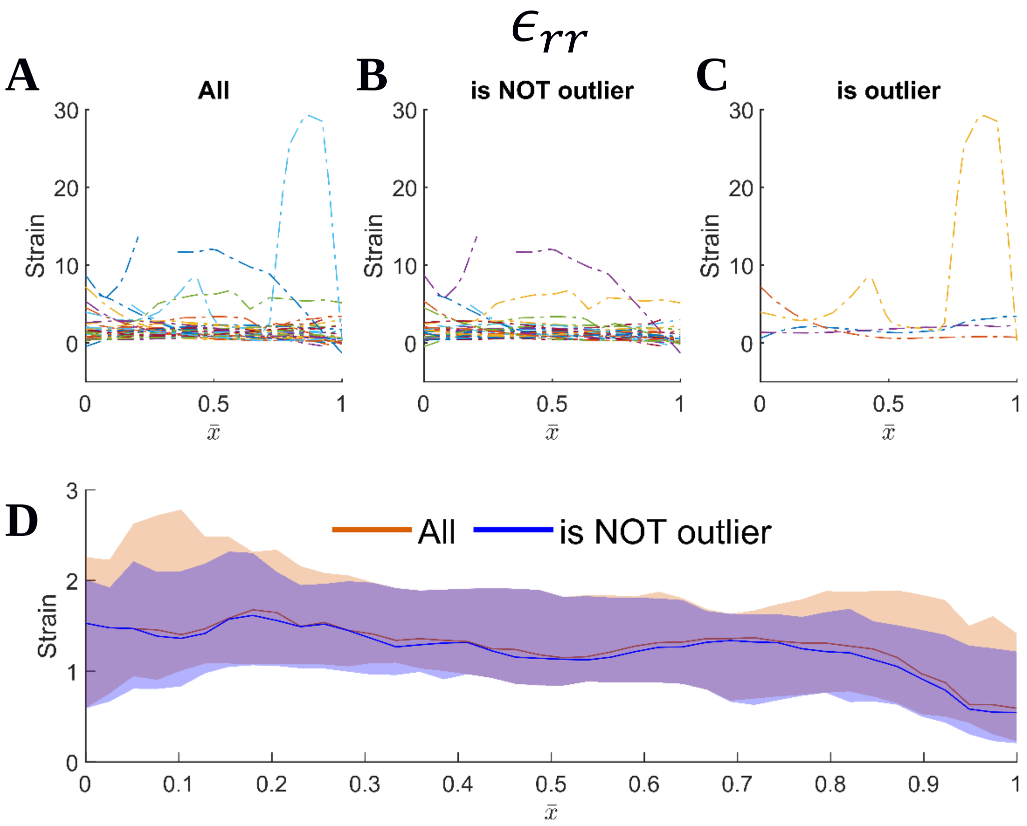


**Figure S5: (A** for each strain component**)** The raw data associated with the strain component measurement from the DIC analyses. We used the Hausdorff Distance [HD (Danziger, 2021)] to help identify outliers. This metric evaluates the similarity of two curves, with a lower HD number indicating higher similarity. We evaluated the distance of the strain components curves from the median response for each strain component and identified the outliers using an automatic outlier detection scheme based on data quartiles, where samples with HD greater than three scaled median absolute deviations (MAD) away from the median were considered outliers (*isoutlier()* function; Matlab). The data set after outlier truncation (**B** for each strain component) and the detected outliers (**C** for each strain component**)**. The median and IQR of strain components before and after truncation of outliers **(D** for each strain component**)**. “All” includes all scans; “is NOT outlier” excludes outliers.

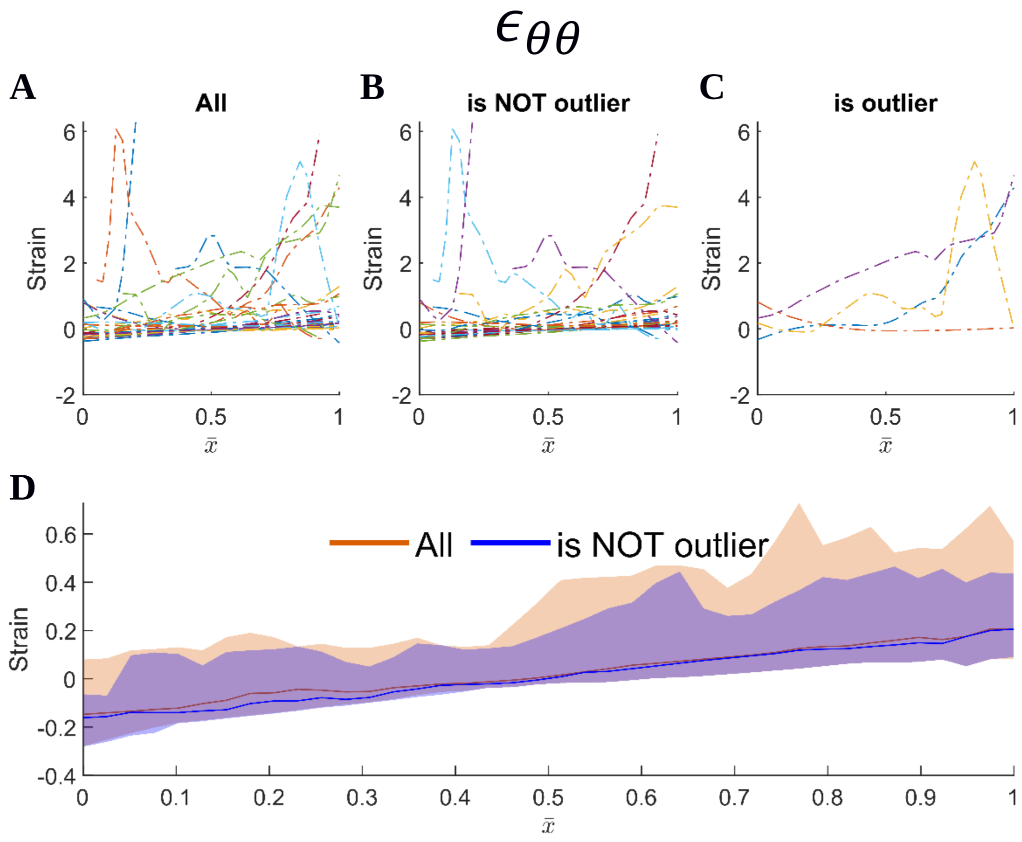


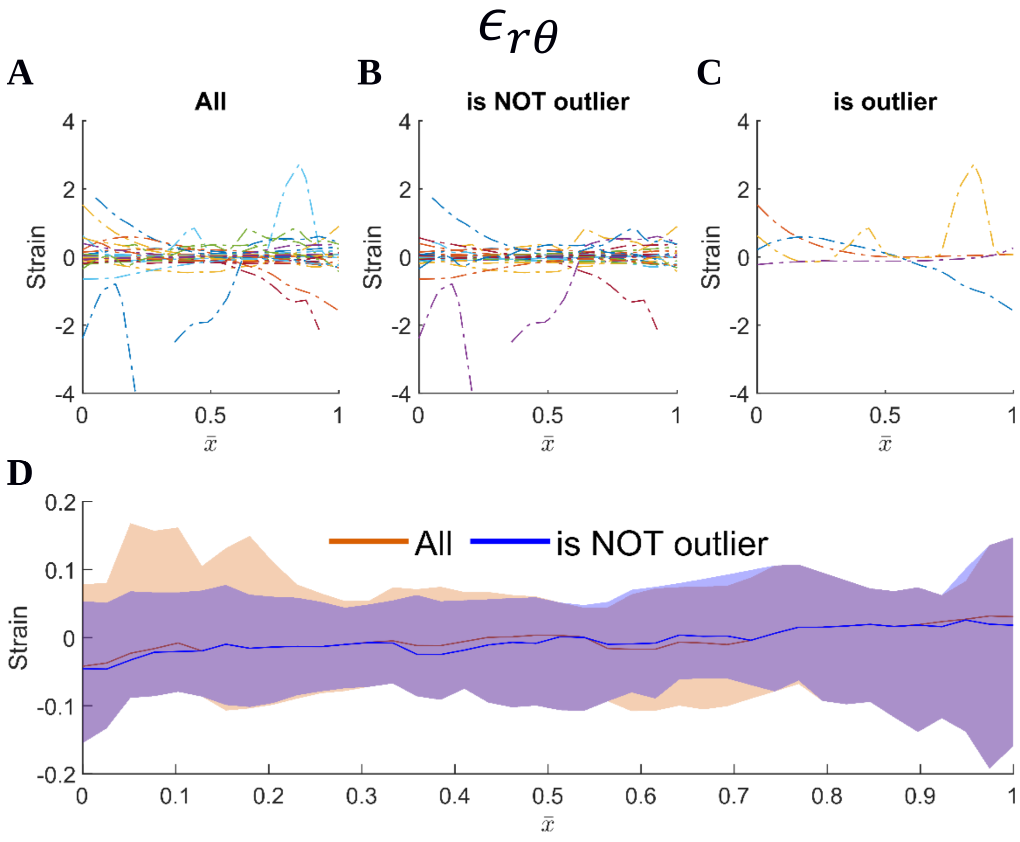


**Figure S5:** continued

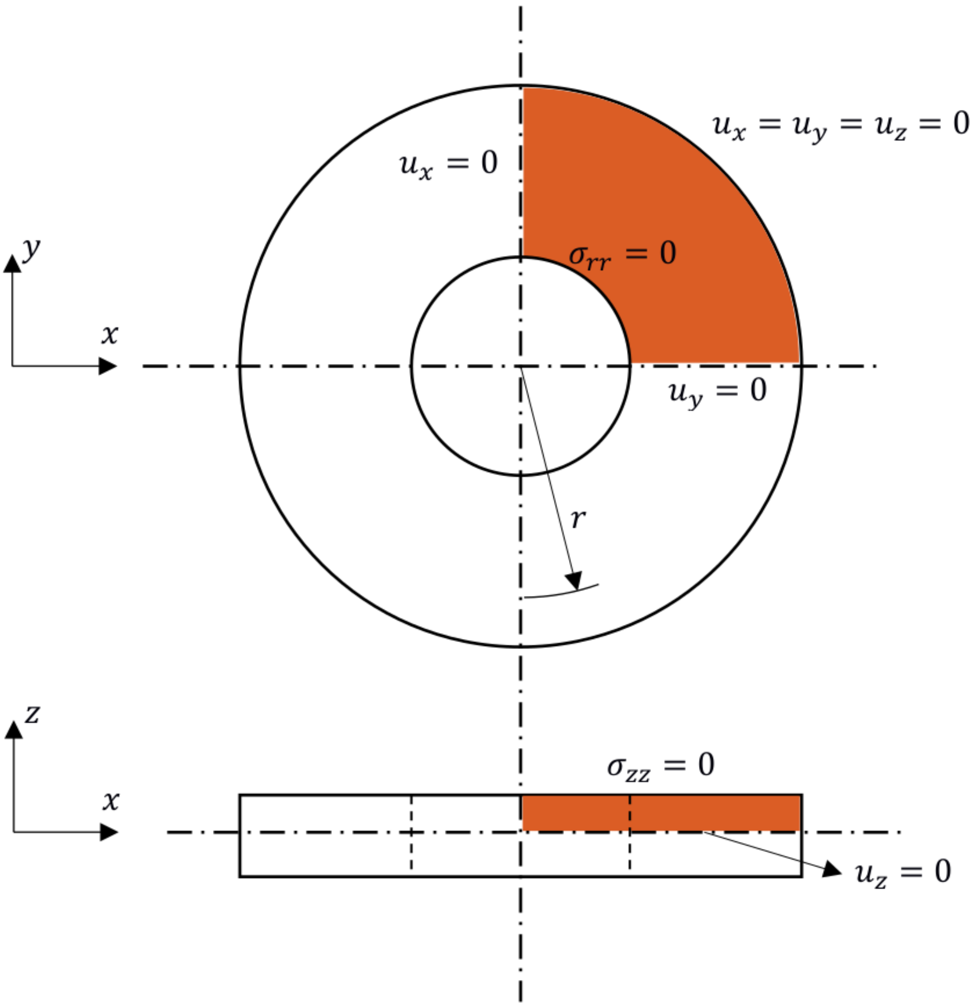


**Figure S6**: The geometry and boundary conditions of the eight-fold symmetric finite element model used in this study.

### References

Hausdorff Distance [WWW Document], n.d. URL https://www.mathworks.com/matlabcentral/fileexchange/26738-hausdorff-distance (accessed 11.4.21).
